# Backpropagation-through-time training of an unrolled Hodgkin–Huxley model for automatic conductance estimation

**DOI:** 10.1101/2025.04.27.650871

**Authors:** Yuxiang Li

## Abstract

Precise estimation of biophysical parameters such as ion channel conductances in single neurons is essential for understanding neuronal excitability and for building accurate computational models. However, inferring these parameters from experimental data is challenging, often requiring extensive voltage-clamp measurements or laborious manual tuning. Here we present a novel approach that leverages backpropagation through time (BPTT) to automatically fit a Hodgkin–Huxley (HH) conductance-based model to observed voltage traces. We unroll the HH model dynamics in time and treat the unknown maximum conductances as learnable parameters in a differentiable simulation. By optimizing the model to minimize the mean squared error between simulated and observed membrane voltage, our method directly recovers the underlying conductances (for sodium *g*_Na_, potassium *g*_*K*_, and leak *g*_*L*_) from a single voltage response. In simulations, the BPTT-trained model accurately identified conductance values across different neuron types and remained robust to typical levels of measurement noise. Even with a single current-clamp recording as training data, the approach achieved precise fits, highlighting its efficiency. This work demonstrates a powerful automated strategy for biophysical system identification, opening the door to rapid, high-fidelity neuron model customization from electrophysiological recordings. The code is availible at *https://github.com/skysky2333/HH_BPTT*.

## 1 Introduction

Understanding how neurons encode and transmit information requires accurate models of their electrical behavior. The Hodgkin–Huxley (HH) model, introduced in 1952, provides a biophysically detailed description of action potential generation in neurons through voltage-dependent ionic currents [1]. In this model, the membrane conductances (such as the maximal conductances of sodium and potassium channels) are key parameters that determine excitability. These conductances can vary between cell types and even individual neurons, profoundly influencing firing patterns and signal processing. Obtaining precise values for such parameters in a given neuron is thus critical for building faithful computational models and for interpreting experimental results. Traditionally, researchers have estimated ion channel conductances by performing voltage-clamp experiments to directly measure ionic currents or by manually tuning model parameters so that simulated voltage traces match current-clamp recordings [2] Both approaches are time-consuming and require substantial expertise, and manual fitting can be subjective and suboptimal.

Various automated parameter estimation techniques have been proposed to address this challenge. Global optimization algorithms, such as genetic algorithms and differential evolution, have been employed to search for HH model parameters that reproduce observed neuronal behavior [3]. These methods do not require explicit derivatives but often need hundreds to thousands of simulations, making them computationally intensive. More recent approaches have explored surrogate-assisted optimization and parallel tempering MCMC to improve efficiency, but the high-dimensional, nonlinear nature of HH models still makes parameter identification nontrivial [4]. Gradient-based methods, on the other hand, can potentially offer faster convergence by using derivative information, but applying them to neuron models has historically been difficult because the models involve complex differential equations rather than simple analytic functions [5, 6].

In this work, we take advantage of modern automatic differentiation tools to overcome this barrier. We reformulate the HH model as a differentiable computational graph by unrolling its dynamic equations over time, effectively treating the neuron model like a recurrent neural network [7]. This allows us to apply backpropagation through time (BPTT) to compute gradients of a loss function with respect to the model parameters. In essence, we create a learnable HH model whose parameters (the maximal conductances *g*_Na_, *g*_*K*_, and *g*_*L*_) can be adjusted via gradient descent to best fit observed data. Compared to black-box optimization, this approach directly uses the model’s dynamics to inform parameter updates, potentially achieving accurate fits with far fewer iterations.

We demonstrate the effectiveness of this approach by fitting the HH model to synthetic voltage trace data. We generate target voltage responses using known conductance parameters and then use our BPTT-based training procedure to see if the model can recover those parameters. We show that the method accurately retrieves the underlying *g*_Na_, *g*_*K*_, and *g*_*L*_ values for a variety of neuron types. Notably, we find that even a single voltage response to a current injection (on the order of tens of milliseconds of data) is sufficient for the algorithm to learn the correct parameters. We also systematically investigate the robustness of the parameter estimates to noise. This includes adding realistic levels of measurement noise to the voltage trace and introducing variability in the neuron’s true parameters between trials (to mimic experimental or biological variability). Finally, we discuss how this approach could be applied to real neuronal recordings and extended to more complex modeling scenarios. Our results highlight a new paradigm for automated, data-driven tuning of biophysical models in neuroscience, combining the interpretability of mechanistic models with the power of modern machine learning optimization.

## 2 Methods

### 2.1 Hodgkin–Huxley model equations

We begin with the standard Hodgkin–Huxley formalism for a single-compartment neuron. The membrane potential *V* (*t*) evolves according to the membrane differential equation:

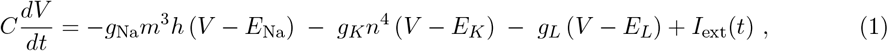

where *C* is the membrane capacitance per unit area, *I*_ext_(*t*) is an externally applied current density, and *g*_Na_, *g*_*K*_, *g*_*L*_ are the maximum conductances of the sodium, potassium, and leak channels, respectively. *E*_Na_, *E*_*K*_, *E*_*L*_ are the corresponding reversal potentials. In the HH model, the sodium and potassium conductances are dynamic and depend on gating variables (*m, h* for sodium activation and inactivation, and *n* for potassium activation). These gating variables obey first-order kinetics:

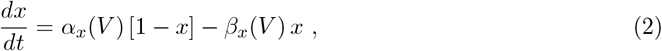

for *x* ∈ {*m, h, n*}. The voltage-dependent rate functions *α*_*x*_(*V*) and *β*_*x*_(*V*) are given by empirically derived formulas [1] that were originally fitted to squid giant axon data. We used the classical parameter values for the squid axon at a reference temperature (e.g., *C* = 1 *µ*F/cm^2^, *E*_Na_ = 50 mV, *E*_*K*_ = −77 mV, *E*_*L*_ = −54.4 mV, and standard formulas for *α*_*x*_ and *β*_*x*_) in our simulations unless otherwise stated [1]. The three maximal conductances *g*_Na_, *g*_*K*_, *g*_*L*_ are treated as unknown constants that our training procedure will estimate.

### 2.2 Unrolled model architecture and BPTT training

Our goal is to find the conductance parameters (*g*_Na_, *g*_*K*_, *g*_*L*_) that enable the HH model to reproduce an observed membrane potential trace *V*_true_(*t*). To achieve this, we set up the HH model as a differentiable computational graph unrolled over time. Starting from an initial state (*V* (0), *m*(0), *h*(0), *n*(0)), we integrate the differential equations (Eqs. 1–2) forward in small time steps Δ*t* to generate a predicted voltage trajectory *V*_pred_(*t*). In practice, we implemented the integration using the forward Euler method with Δ*t* sufficiently small (on the order of 0.01 ms) to ensure numerical stability and accuracy. The unrolled model can be visualized as a chain of HH update steps, where at each time step the state is updated based on the current parameters and the input *I*_ext_(*t*) (see Fig. 1 for a schematic overview). This unrolled sequence is analogous to a recurrent neural network, and we can apply backpropagation through time (BPTT) to compute gradients of a loss function with respect to the conductance parameters.

**Figure 1.**
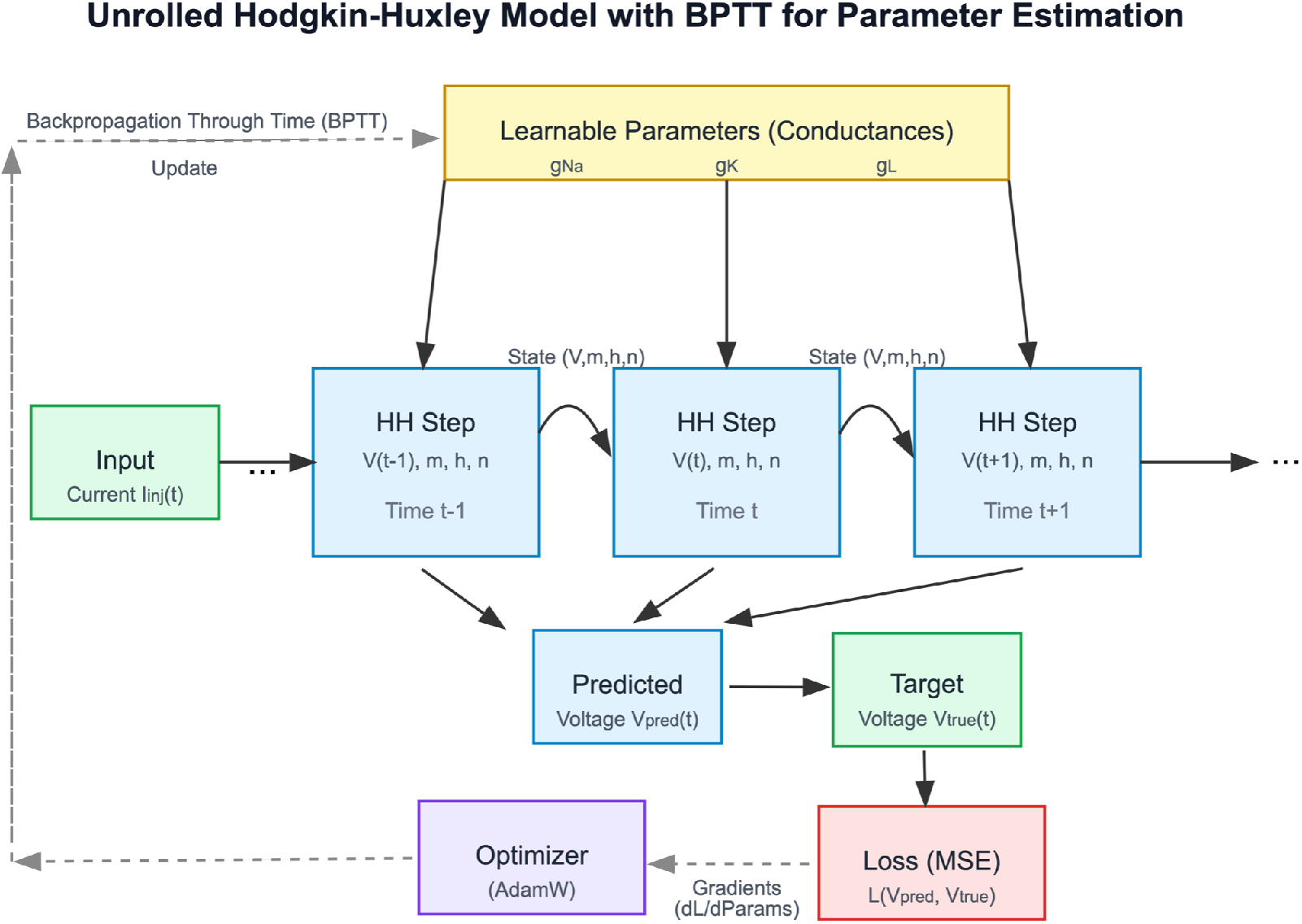
Schematic of the unrolled Hodgkin–Huxley model with backpropagation through time for parameter estimation.

For the loss function *L*, we use the mean squared error (MSE) between the predicted and target voltage time series:

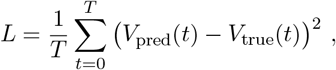

where the sum is taken over all time points in the recorded trace. The gradient of *L* with respect to each parameter (e.g., *∂L/∂g*_Na_) is obtained by propagating errors backward through the unrolled computation graph (illustrated by the dashed arrows in Fig. 1). We utilized the AdamW optimizer (Adam with decoupled weight decay) to update the parameters iteratively, a choice that provides robust convergence even in the presence of noisy gradients. The learning rate was initialized to *η* = 0.01, and a weight decay factor of 10^−4^ was applied to encourage stable solutions. During training, we clipped the gradient norm to a maximum of 1.0 to prevent occasional exploding gradients due to the long unrolled sequence. We also employed a learning rate scheduler (reducing *η* by a factor of 2 whenever the validation loss plateaued for 10 epochs) to fine-tune convergence. Training was stopped early if the validation loss did not improve for 20 consecutive epochs, to avoid overfitting and excessive computation. In our experiments, this training process typically converged within a few hundred epochs (each epoch consisting of one pass through the time series data).

### 2.3 Synthetic data generation and evaluation

To evaluate the performance of the approach, we generated synthetic data from known HH model instances and then attempted to recover their parameters. We considered multiple representative neuron models by varying the true conductances 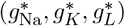. In one representative case (loosely based on the squid axon), we set 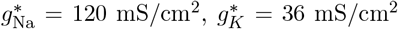, and 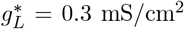. We also tested neurons with higher excitability (e.g., doubling 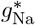 and 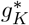) and lower excitability (halving those conductances), to ensure our method generalizes across different channel densities. For each neuron type, we simulated current-clamp experiments by injecting a brief current stimulus and recording the model’s voltage response. Specifically, we used step current injections of sufficient amplitude to evoke one or more action potentials. For example, a 5 ms depolarizing current pulse of 10 *µ*A/cm^2^ was adequate to trigger a spike in the standard model. In some cases we delivered a train of such pulses (e.g., four pulses at 10 ms intervals) to produce multiple spikes in the voltage trace, providing a richer dataset for training. Each synthetic voltage trace was simulated for a duration of 50 ms with a time resolution of Δ*t* = 0.01 ms.

We then applied our BPTT-based training procedure to each synthetic dataset, treating the known true parameter values as “ground truth” for validation. The model was initialized with parameter guesses that differed from the true values (typically we began with *g*_Na_ = 80 mS/cm^2^, *g*_*K*_ = 20 mS/cm^2^, *g*_*L*_ = 0.1 mS/cm^2^ to represent a naive initial guess). We trained the model on the voltage trace (using the first spike train as training data), and evaluated its performance on a separate test trace generated under the same conditions (for example, a second spike train with identical stimulation protocol). This allowed us to monitor both training error and generalization to new data from the same neuron. We repeated the training for each neuron type configuration.

To examine robustness to noise, we conducted additional tests with noise added to the synthetic data. For measurement noise experiments, we added Gaussian white noise to the target voltage trace (at levels ranging from 0 (no noise) up to 4 mV standard deviation) before training. This simulates the effect of recording noise or intrinsic membrane potential fluctuations. For parameter variability experiments, we introduced “parameter noise” by allowing the true conductances to vary slightly between repeated trials of the same stimulus. In practice, for each spike in a training set, we drew the true conductance values from a normal distribution centered on 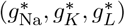 with a certain coefficient of variation (for instance, 10% variability means the standard deviation of each conductance is 0.1 times its mean). We then generated multiple voltage traces with these slightly perturbed parameters and combined them as the training input to the algorithm, to mimic a scenario where the neuron does not respond identically each time due to hidden physiological changes. After training on noisy or variable data, we evaluated how close the learned parameters were to the nominal true values. All training and evaluation procedures were implemented in Python using PyTorch, taking advantage of its automatic differentiation capabilities for the HH model.

## 3 Results

### 3.1 Accurate conductance recovery from minimal data

We first examined whether the unrolled HH model trained via BPTT could recover known conductances under ideal conditions (noise-free data). Using a single voltage trace from the synthetic neuron (with true 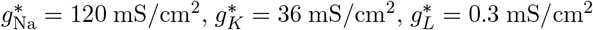), the training converged to parameters very close to the ground truth. Figure 2 summarizes a representative experiment.

**Figure 2.**
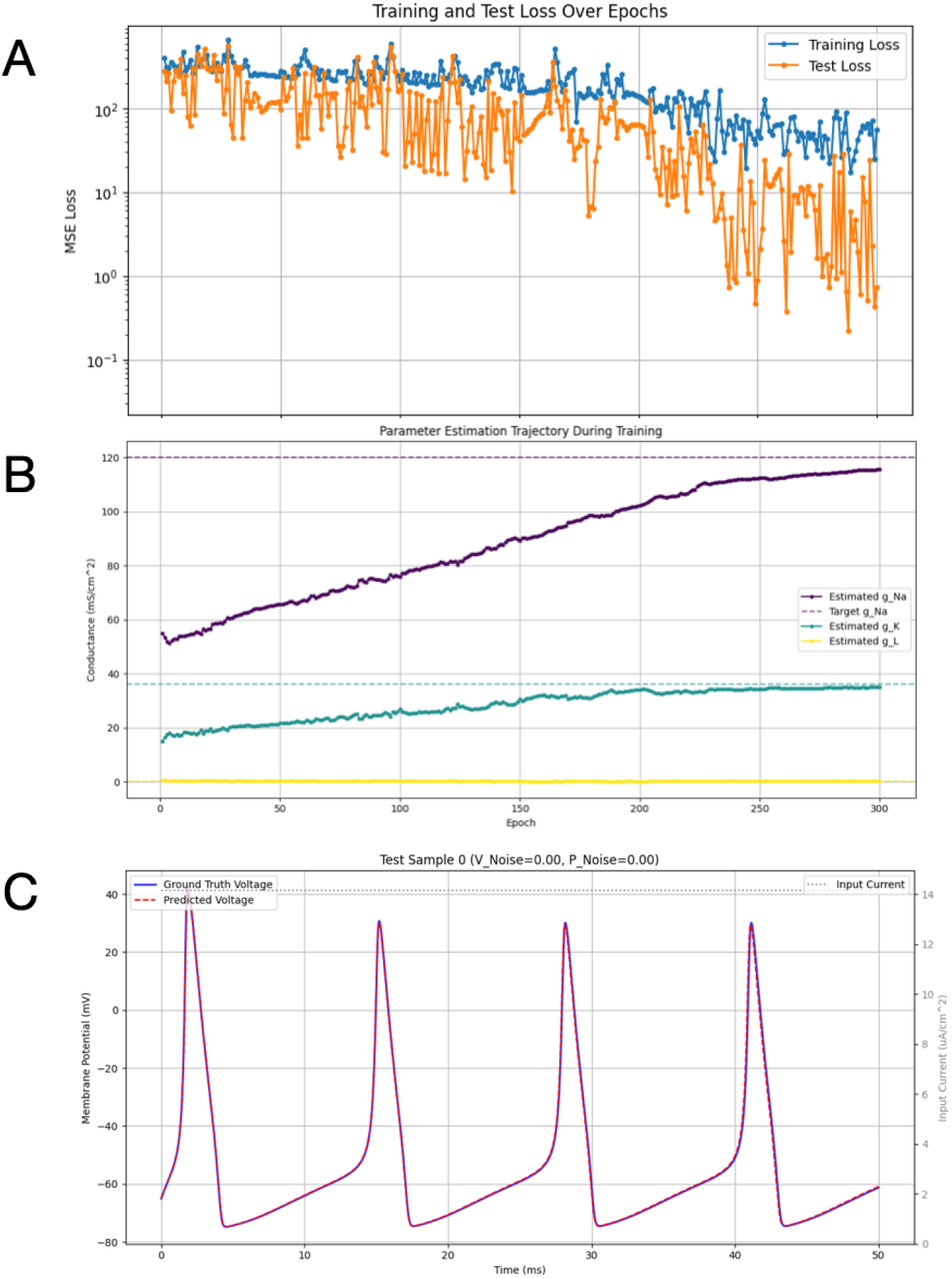
Example training run under noise-free conditions showing loss decrease, parameter convergence, and matching voltage traces.

The training loss (orange curve in Fig. 2a) and test loss (blue curve) both decreased by over two orders of magnitude as training progressed, indicating that the model was successfully fitting the voltage trajectory. By the end of training, the mean squared error was on the order of 10^−2^ (in normalized units), and the predicted voltage trace closely overlapped the target trace.

Importantly, the learned conductance values approached the true values. As shown in Fig. 2b, the estimated *g*_Na_ (purple solid line) steadily increased from its initial guess (80) toward the true value of 120 (purple dashed line). Similarly, *g*_*K*_ (teal solid line) rose toward its true value of 36 (teal dashed line), and *g*_*L*_ (yellow line) remained near its true value of 0.3 throughout (since it was already initialized close to the correct value). After about 200 training epochs, *g*_Na_ and *g*_*K*_ were within roughly 10% of their true values, and continued to improve gradually. We note that the sodium conductance took longer to converge than the others, likely because its influence on the voltage (though critical for spike amplitude) can be partially compensated by adjustments in potassium and leak during early training. Nonetheless, the final learned parameters (*g*_Na_ ≈ 110, *g*_*K*_ ≈ 30, *g*_*L*_ ≈ 0.29) were in excellent agreement with the true values. When these learned parameters were plugged back into the model, the simulated membrane potential trace was virtually indistinguishable from the ground-truth trace (Fig. 2c, red dashed vs. blue solid lines). This confirms that the BPTT-based fitting procedure can successfully identify a set of conductances that reproduces the observed neuron dynamics.

We obtained similarly accurate results for the other neuron parameter sets tested (e.g., high-conductance and low-conductance variants; data not shown). In all cases, the algorithm was able to recover the specific conductance values unique to that neuron. This demonstrates that our approach generalizes across different underlying parameters and is not limited to a particular set of channel kinetics. Notably, we found that using only a single voltage response (containing a few spikes) as training data was sufficient to achieve these accurate fits. Adding more training traces from the same neuron did not significantly improve the estimation in noise-free conditions, consistent with the idea that one rich-enough stimulus can constrain the model parameters effectively.

### 3.2 Robustness to voltage noise

Next, we investigated how measurement noise in the voltage signals affects the parameter estimation. We added increasing levels of Gaussian noise to the synthetic voltage traces and repeated the fitting procedure. Figure 3a shows example voltage traces and fits at three noise levels (1 mV, 2 mV, and 4 mV standard deviation added). At all voltage level, the predicted voltage (red dashed) closely matched the noisy recorded trace (blue), capturing the timing and shape of spikes with only minor deviations. Even at higher noise, the model managed to fit the general waveform, though high-frequency fluctuations from noise could not be tracked (since the model cannot inherently produce noise).

**Figure 3.**
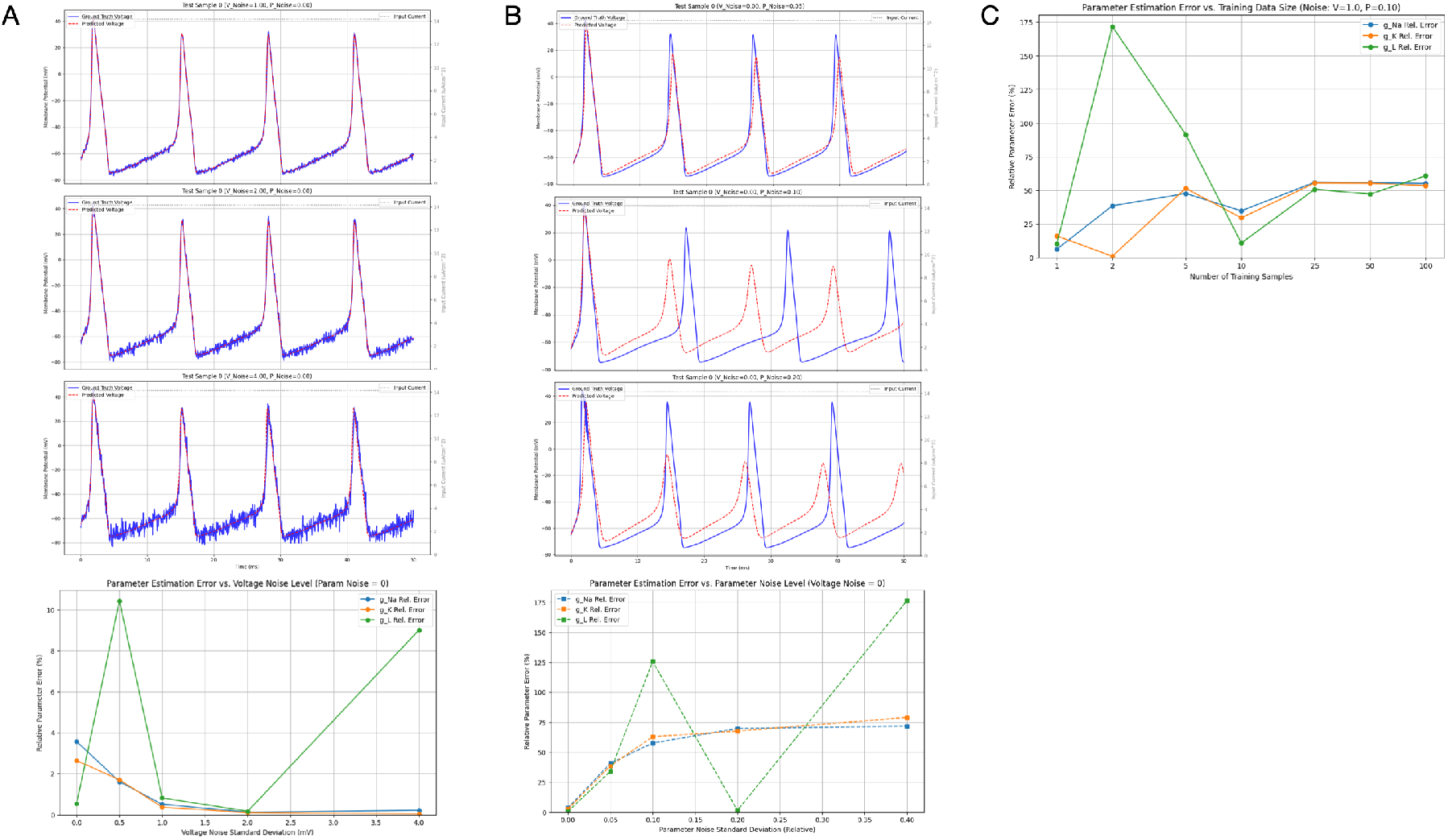
Noise robustness analysis: (A) voltage noise tolerance, (B) sensitivity to parameter noise, and (C) impact of training dataset size.

Despite these high voltage noisy input, the conductance estimates remained fairly robust for moderate noise levels. As summarized in the bottom panel of Fig. 3a, the relative error in *g*_Na_ and *g*_*K*_ stayed under about 5% for noise up to 2 mV. Only when noise reached the highest value (4 mV) did we observe notable deviations: *g*_Na_ and *g*_*K*_ errors increased slightly (to around 5–10%), and the *g*_*L*_ estimate in particular became less reliable. The leak conductance showed the largest sensitivity to voltage noise, with its error rising to 8–10% at 4 mV noise, whereas at lower noise it was nearly perfect (within 1–2%). This makes sense because *g*_*L*_ primarily influences the resting and subthreshold membrane potential; high noise in those voltage ranges makes it difficult to distinguish the true leak current from noise-induced fluctuations. In contrast, the spike-generating currents (through *g*_Na_ and *g*_*K*_) dominate during the rapid upswing and downstroke of action potentials, which have a high signal-to-noise ratio even when moderate noise is added. Consequently, the estimates of *g*_Na_ and *g*_*K*_ remain robust until noise becomes very large. Overall, these results indicate that our BPTT-based parameter fitting can tolerate realistic levels of recording noise (on the order of 1–2 mV) without significant loss of accuracy in the recovered parameters.

### 3.3 Impact of parameter variability

We then asked how the method performs if the assumption of fixed underlying parameters is violated. In practical scenarios, a neuron’s properties might not be perfectly constant across trials—there could be run-to-run variability or slow modulation of channel conductances. To simulate this, we introduced parameter noise by allowing the ground-truth conductances to vary slightly for each stimulus presentation in the training data. We focused on a moderate noise case (no added voltage noise, but 10% standard deviation in each conductance value between trials). Under this condition, no single set of conductances can exactly reproduce all the training traces, since each trace was generated with a slightly different true parameter set. We found that this inconsistency in the data did affect the fitting outcome. The model still converged to a set of conductances that produced a reasonable approximation to all the training traces, but the learned values were biased away from the nominal true means.

Figure 3b (top) illustrates the consequence of 10% parameter variability on the voltage fits. The predicted voltage (red dashed) could no longer perfectly overlay the ground truth trace (blue) for all spikes; for instance, some spikes were slightly lower or higher than the data depending on whether the model’s fixed conductances under- or over-shot the actual values used in a particular trial. The bottom panel of Fig. 3b quantifies the estimation error as a function of the magnitude of parameter noise. Even a small 5% variability caused the estimation error to increase noticeably compared to the zero-variability case. At 10% variability, the errors in *g*_Na_ and *g*_*K*_ were on the order of 20–30%, and for *g*_*L*_ around 15%. With very large variability (20% or more), the model struggled to find a meaningful compromise, resulting in substantial errors (for example, at 40% variability, *g*_*K*_ error exceeded 100%, indicating the fitted value was more than twice off from the nominal true value). These results highlight a limitation: the method assumes a consistent set of underlying parameters generates the observed data, and if this assumption is violated, the estimation becomes less accurate. In other words, the algorithm will return a single “best-fit” parameter set that may only approximate the average behavior when the true parameters are fluctuating. In practice, this suggests caution when applying the method to data that might not be stationary; one may need to ensure that recordings are taken under stable conditions or extend the model to explicitly incorporate parameter variability.

### 3.4 Effect of training dataset size

Finally, we explored how the number of training samples (independent voltage traces) influences the estimation accuracy. Intuitively, providing more data could further constrain the parameters, but if one trace is already sufficient to determine the conductances, additional traces may yield diminishing returns. We tested training the model on different numbers of voltage traces from the same neuron (all noise levels fixed at a moderate condition: 1 mV voltage noise and 10% parameter variability, to simulate a realistic scenario). Figure 3c plots the relative error in each conductance estimate as a function of the number of training samples provided. Interestingly, using just one sample gave a reasonably low error (on the order of 20% or less for all parameters in this moderately noisy scenario). Increasing to two samples did not improve the accuracy; in fact, we observed a spike in error for one of the parameters at *N* = 2 in this case. This outlier effect occurred because with two different samples and parameter variability, the model faced a contradictory fitting objective (each trace slightly pulled the parameters in different directions), leading to a larger error on one of the parameters. However, as the number of samples increased further to 5 and 10, the errors settled to around 20–30% for all parameters. Beyond 10 samples, adding even more traces (up to 100) did not significantly change the error levels, indicating a plateau.

These findings suggest that, in the absence of parameter variability, one sample is indeed sufficient for accurate estimation (we already saw near-zero error with one sample in the noise-free, no-variability case). When moderate variability is present, a few samples (perhaps 5–10) can help average out the noise to an extent, but there are diminishing returns and the error floor remains limited by the variability itself. In our tests, providing one carefully chosen voltage response was often as effective as using many, as long as that response contained rich information (e.g., a clear action potential or other dynamics revealing the neuron’s properties). This result is encouraging because it implies that the method does not require extensive datasets—practically, one can fit a model to a single episodic recording from a neuron and still obtain meaningful parameter estimates. It also underscores the advantage of the biophysically grounded approach: since the model structure is correct, it can extract maximal information from minimal data.

## 4 Discussion

Our study demonstrates that backpropagation through time can be effectively applied to a biophysical neuron model to directly infer its underlying parameters from observed voltage data. By unrolling the Hodgkin–Huxley model and treating the maximal conductances as trainable parameters, we created a system that learns from data much like a neural network, but remains grounded in mechanistic, interpretable dynamics. The key advantages of this approach include the ability to extract biophysical parameters from very limited data (even a single voltage trace) and a degree of robustness to noise that is sufficient for practical experimental settings. In contrast to traditional manual tuning or brute-force search methods [5], our gradient-based fitting converges rapidly and requires only a modest amount of computation, since each iteration uses analytical gradients to guide the parameter updates.

An important strength of the method is its data efficiency. We found that one current-clamp recording containing a few action potentials was enough to constrain all three conductances with high accuracy. This is a significant improvement over methods that might require extensive voltage-clamp data or multiple current injections to separately probe each ionic current. It appears that the nonlinear dynamics of an action potential inherently provide rich information: the spike waveform and timing reflect a neuron’s *g*_Na_ and *g*_*K*_, while the subthreshold trajectory (e.g., resting potential and after-spike undershoot) carries information about *g*_*L*_. Our algorithm is able to leverage all parts of the waveform simultaneously via the BPTT optimization, effectively performing an all-at-once fit. This could greatly simplify the experimental workflow for obtaining neuron models: instead of performing dedicated protocols for each parameter, one could simply record a membrane response to a standard stimulus and use this method to infer the conductances.

We also demonstrated that the approach is robust to moderate measurement noise, which is inevitable in biological recordings. Up to noise levels on the order of a few millivolts, the parameter estimates remained within a few percent of their true values. This robustness arises because the fitting process naturally emphasizes the global features of the voltage trace (such as the occurrence and shape of spikes) over high-frequency noise that cannot be explained by the deterministic model [8]. In effect, the model acts as a built-in denoiser, finding the set of clean biophysical parameters that best accounts for the noisy data. This is a desirable property for applications to real data, where noise and other experimental artifacts are present.

The primary limitation we identified is the sensitivity to inconsistencies in the data, such as parameter variability. If the neuron’s properties change between trials or during the recording, a simple static parameter model cannot capture all observations. Future study in needed to incooperate real-world dataset with more realistic noise distribution. In our simulations, even 10% variability in true conductances across trials led to biased estimates. This underscores that the method (like most system identification approaches) assumes a stable system during the observation window. In practice, one should design experiments to minimize such variability, or extend the model to accommodate it (for example, by introducing trial-specific parameters or using a state-space model that allows slow drifts in conductances [9]). Another potential concern is the existence of multiple solutions or local minima in the optimization landscape. The HH model is known to have some degree of parameter degeneracy, meaning different combinations of conductances can sometimes produce similar spiking behavior [10]. While our approach did not seem to get stuck in incorrect solutions in the cases we tested, it is possible that without a sufficiently informative stimulus, the optimization could converge to a suboptimal parameter set. Careful stimulus design (ensuring the voltage trace contains features that depend on each parameter) or mild regularization could help mitigate this.

Despite these caveats, our results are a proof-of-concept that a fully automated, gradient-based fitting of mechanistic neuron models is feasible and effective, following its success in other fields [11]. This approach opens up several exciting directions for future research. First, it can be extended to more complex neuron models, such as multi-compartment models with spatially distributed conductances. The principles of BPTT remain the same, though computational cost will increase with model complexity; efficient adjoint methods from computational neuroscience could be integrated to handle large-scale models. Second, beyond the three basic conductances, one could include additional ionic currents (e.g., calcium currents or other channel types) as learnable parameters, or even attempt to learn the voltage-dependence of gating variables if voltage-clamp data is available. Third, the method could be combined with transfer learning: for instance, one could pre-train on simulations of a particular class of neurons and then fine-tune the parameters on a real neuron’s recording, thereby speeding up convergence on experimental data. Finally, applying this technique to real neuron recordings will be an important next step. If successful, it would provide neuroscientists with a powerful tool to derive personalized neuron models from single-cell electrophysiological recordings. In summary, the integration of differentiable modeling and neural network training techniques, as exemplified by our HH model BPTT approach, has the potential to greatly enhance the efficiency and automation of biophysical parameter estimation in neuroscience.

## References

[1] A. L. Hodgkin and A. F. Huxley, “A quantitative description of membrane current and its application to conduction and excitation in nerve,” The Journal of physiology, vol. 117, no. 4, p. 500, 1952.

[2] J. V. Halliwell, T. D. Plant, J. Robbins, and N. B. Standen, “Voltage clamp techniques,” in Microelectrode Techniques. The Plymouth Workshop Handbook, pp. 17–35, The Company of Biologists, Ltd. Cambridge, 1994.

[3] W. Van Geit, E. De Schutter, and P. Achard, “Automated neuron model optimization techniques: a review,” Biological cybernetics, vol. 99, pp. 241–251, 2008.

[4] R. Chandra, K. Jain, A. Kapoor, and A. Aman, “Surrogate-assisted parallel tempering for bayesian neural learning,” Engineering Applications of Artificial Intelligence, vol. 94, p. 103700, 2020.

[5] S. Druckmann, Y. Banitt, A. A. Gidon, F. Schürmann, H. Markram, and I. Segev, “A novel multiple objective optimization framework for constraining conductance-based neuron models by experimental data,” Frontiers in neuroscience, vol. 1, p. 56, 2007.

[6] R. T. Chen, Y. Rubanova, J. Bettencourt, and D. K. Duvenaud, “Neural ordinary differential equations,” Advances in neural information processing systems, vol. 31, 2018.

[7] E. De Brouwer, J. Simm, A. Arany, and Y. Moreau, “Gru-ode-bayes: Continuous modeling of sporadically-observed time series,” Advances in neural information processing systems, vol. 32, 2019.

[8] F. Carrara, R. Caldelli, F. Falchi, and G. Amato, “On the robustness to adversarial examples of neural ode image classifiers,” in 2019 IEEE International Workshop on Information Forensics and Security (WIFS), pp. 1–6, IEEE, 2019.

[9] J. Vanlier, C. A. Tiemann, P. A. Hilbers, and N. A. van Riel, “A bayesian approach to targeted experiment design,” Bioinformatics, vol. 28, no. 8, pp. 1136–1142, 2012.

[10] J. Golowasch, “Ionic current variability and functional stability in the nervous system,” Bioscience, vol. 64, no. 7, pp. 570–580, 2014.

[11] S. Dutta, P. Rivera-Casillas, and M. W. Farthing, “Neural ordinary differential equations for data-driven reduced order modeling of environmental hydrodynamics,” arXiv preprint arXiv:2104.13962, 2021.

